# Methane and ethane production rates by methyl-coenzyme M reductase in cell extracts from different methanogens

**DOI:** 10.1101/2024.01.08.574666

**Authors:** V. P. Thinh Nguyen, Sebastian Keller, Silvan Scheller

## Abstract

The enzyme methyl-coenzyme M reductase (Mcr) is responsible for most of the biologically produced methane. Mcr catalyzes the reversible conversion of methyl-coenzyme M and coenzyme B to methane and the corresponding heterodisulfide CoM-S-S-CoB as the last step in all methanogens. In anaerobic methanotrophs, it catalyzes the same reaction to proceed in the reverse direction.

While Mcr has been historically considered as an enzyme exclusive to the one-carbon metabolism, recent studies have demonstrated that homologs of Mcr have evolved to catalyze the first step of the archaeal anaerobic oxidation of medium and long-chain alkanes to their corresponding alkyl-coenzyme M thioethers.

While ample metagenomic studies and data related to Mcr is available, in vitro experiments are limited, because purifying Mcr in its active nickel-I form is challenging. Fully active enzyme has only been reported for *Methanothermobacter marburgensis* isoenzyme I, from which most of the enzyme studies have been performed.

To get an overview of the substrate scope of different Mcr variants, we tested cell-free lysates from the methanogens *Methanosarcina mazei, Methanosarcina acetivorans, Methanococcus maripaludis, Methanothermococcus okinawensis*, and *M. marburgensis* for their rates to convert methyl- and ethyl-coenzyme M to methane and to ethane, respectively.

Cell extracts from *M. mazei* showed an ethane production rate of ca. 8% relative to methane production at 37 °C, and about 14% at 49 °C in an assay relying on titanium (III) citrate and cobalamin to regenerate coenzyme B from the CoM-S-S-CoB heterodisulfide.

For cell extracts of *M. marburgensis*, we found an ethane-to-methane production rate of 8%, which is substantially higher than the reported value of ca. 0.3% for the ratio of their maximal catalytic rate for purified isoenzyme I. Since *M. marburgensis* is known to express two isoenzymes depending on the growth conditions, we hypothesize that isoenzyme II is substantially more promiscuous towards ethane formation than the well-described isoenzyme I.

Which it is still challenging to obtain accurate kinetic parameters of the Mcr-catalyzed reaction, our experiments demonstrate that Mcr activity can be quickly and conveniently studied *via* cell-free lysates, and that substrate promiscuity towards ethane formation is generally larger than anticipated.

## Introduction

Methanogenic archaea play an important role in the global carbon cycle by producing an estimated global flux of ca. 1 Gt per year of methane [1]. The enzyme responsible for the last catabolic step is methyl-coenzyme M reductase (Mcr) that converts methyl-coenzyme M (Me-S-CoM) and the thiol coenzyme B (CoB-SH) reversibly to methane and the corresponding heterodisulfide (CoB-S-S-CoM) [2] (**Figure 1**). In anaerobic methanotrophic archaea, homologs of Mcr catalyze the same reaction in the reverse direction to activate methane for further catabolism to CO_2_ [3–5]. Recent studies have demonstrated that divergent Mcr homologues have evolved to catalyze the first step of archaeal anaerobic oxidation of medium and short-chain alkanes [6,7].

**Figure 1.**
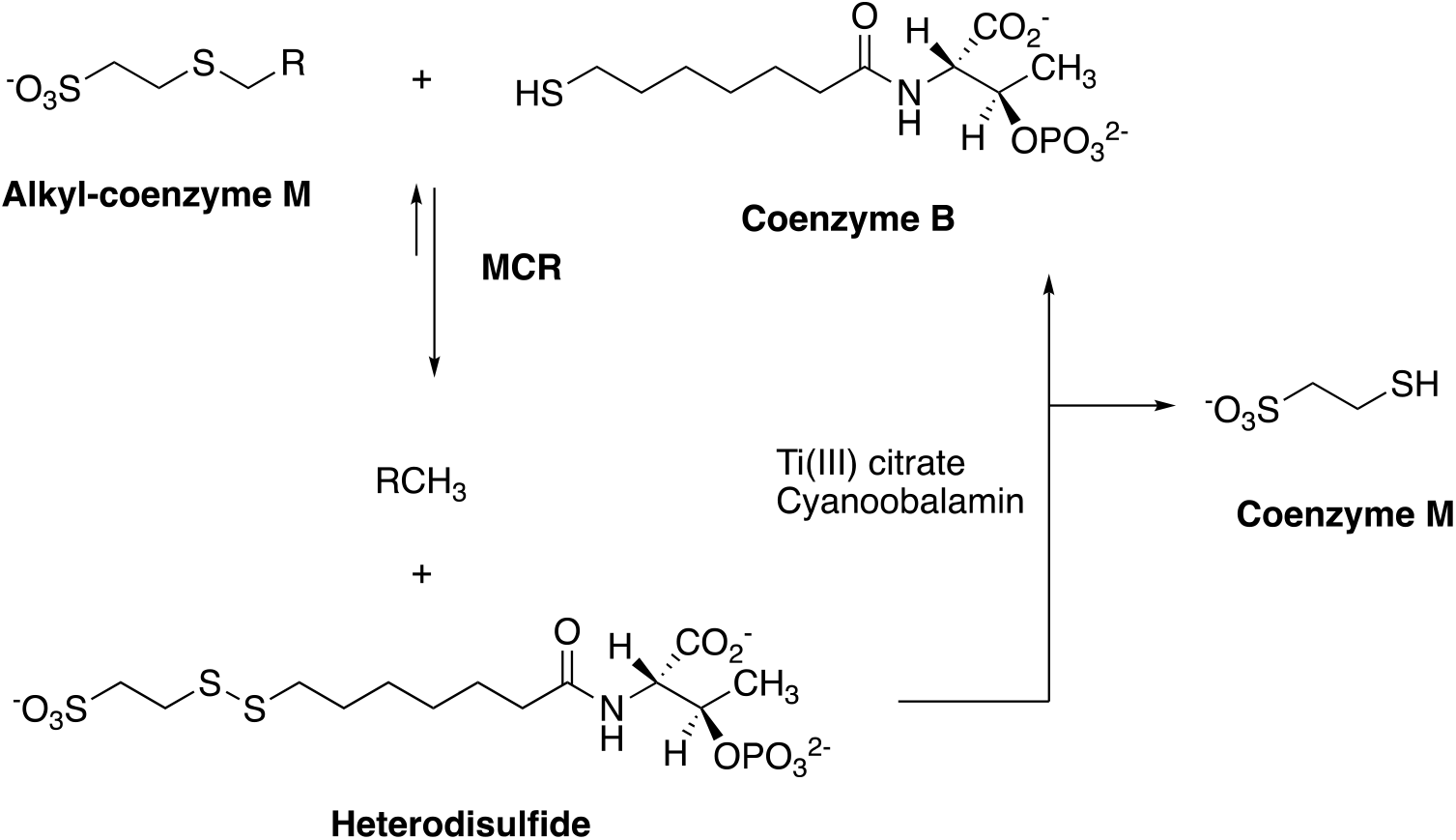
Methyl-coenzyme M reductase-catalyzed conversion of alkyl-coenzyme M to their corresponding alkanes and to the heterodisulfide between coenzymes M and B. The assay includes an excess of Ti(III) citrate and a catalytic amount of cyanocobalamine in order to cleave the heterodisulfide and re-generate coenzyme B. (R = H, CH_3_, C_2_H_5_)

The structure of Mcr holo-enzyme entails three different protein chains in a C_2_-symmetric α_2_β_2_γ_2_ assembly with two active sites, each containing one molecule of the nickel hydrocorphinate F430 as the prosthetic group [8]. Heterologous expression of Mcr is challenging since it contains several post-translational modifications [2,9] that need to be correctly installed, and F_430_ needs to be inserted correctly. In the model methanogen *M. maripaludis*, holo-Mcr from the closely related methanogen *M. okinawensis* could be heterologously expressed with all post-translational modifications correctly installed, and with both cofactors F430 inserted [10]. The enzyme, however, could not be obtained in the active form, which requires its nickel-porphinoid [11] cofactor F_430_ to be in its Ni(I) oxidation state to be active [12,13].

To study the biochemistry of Mcr, pure and fully active Mcr isoenzyme I from *M marburgensis*, which contains two different isoenzymes, have been isolated from 10L bioreactor cultures via strictly anaerobic procedures [14]. Purification of active Mcr from other methanogens appears to be challenging and has not been reported. Traces of oxygen immediately inactivate Mcr, and Mcr is assumed to undergo auto-inactivation during the catalytic cycle. The native activation complex that allows the cells to reduce F430 into the Ni(I) form after its biosynthesis and when deactivated during the catalytic cycle remains to be elucidated. To obtain reliable kinetic *in vitro* data, an enzyme assay containing Ti(III)-citrate and a catalytic amount of cobalamin is required in order to reduce the accumulating heterodisulfide to the co-substrate coenzyme B and generate the weaker inhibitor coenzyme M as a side product (**Figure 1**).

The specific activity of fully active Mcr-I isoenzyme from *M. marburgensis* is reported to be 22.3 U/mg [14]. One study reports a higher value, where the ox1 state has been activated with Ti(III) citrate to red1, a maximal activity of 100 U/mg has been reported [13]. The substrate promiscuity of Mcr-I towards producing ethane from ethyl-S-CoM has been reported to have a relative maximum rate of ethane-to-methane formation equals ca. 0.3% [15]. In a later study that relied on ^13^C-isotope exchange, it could be demonstrated that the rate of C-S bond cleavage is equal for methyl- and for ethyl-coenzyme M, but the ethane stays trapped in the active site and prefers to react back to ethyl-coenzyme M rather than being release as free ethane [16]. The reverse reaction, the conversion of free ethane to ethyl-coenzyme M has been found to occur at a rate of ca. 6% relative to its corresponding C1 reaction, as quantified in a competitive experiment where equimolar pressures of ethane and methane have been incubated with Mcr-I from *M. marburgensis* [17].

Reports of Mcr-catalyzed ethane formation in other methanogen include the ethyl-S-CoM to ethane by cell-free extracts from *Methanosarcina barkeri* under hydrogen supply [18], but no quantification of substrate promiscuity is available from this study. For living cells, different *M. barkeri* strains have been reported to convert mixtures of methanol and ethanol to their corresponding alkanes, with a maximum ethane-to-methane production rate of 0.1% [19]. The conversion of (m)ethanol to (m)ethane putatively involves the two enzymatic steps of methanol:coenzyme M methyl-transferase and Mcr, which makes it impossible to attribute a substrate promiscuity to Mcr.

To estimate the substrate promiscuity of different Mcr variants, we report *in vitro* activity measurements with cell extracts as a simple strategy to rapidly assay the Mcr activity, without the need to purify and reactivate the enzyme. We have adapted a previously published enzyme activity assay of Mcr purified from *M. marburgensis* [8] and optimized the conditions for Mcr-catalyzed ethane formation with cell-free lysate isolated from *M. acetivorans* (strain WWM73) grown on methanol (MeOH). For *M. marburgensis*, we tested the substrate promiscuity depending on growth conditions, and we tested the relative ethane to methane production rate in competitive experiments where mixtures of C1 and C2 substrates are provided.

## Results

The conventional assay (**A**) to measure the methane and ethane formation rate consisted of a master mix that includes 5 mM alkyl-S-CoM, 0.5 mM CoB-homodisulfide, 0.2 mM cobalamin (cyanocobalamin), 20 mM of Ti(III) citrate in 50 mM Tris-HCl at pH = 7.0. The reaction was started by adding 100 µg of cell-free lysate at pH 7 (300 - 600 uL) at 25 °C, with the total volume was kept at 2 mL. Methane and ethane formation were then determined via gas-chromatography (GC-FID).

In addition, we have optimized an assay for high *in vitro* methane and ethane formation using an experimental design approach employing the response surface methodology [20] (**Supporting Information)**. This assay was developed using cell-free lysates from *M. acetivorans* (strain WWM73) grown on methanol (MeOH) [21,22]. The culture was supplemented with 100 mM MeOH and 20 μM sodium sulfide as sulfur source under strict anaerobic conditions. Once the desired growth phase was reached, the cell-free lysate was then prepared from the active culture and will be used to catalyze the conversion of methyl- and ethyl-S-CoM to corresponding methane and ethane, respectively. The optimized assay conditions (**B**) were: OD_600_ at harvesting at moderate/low, 49 °C and 1.1 mM B_12_.

### Testing cell-free lysates from different methanogens for their Mcr-catalyzed methane and ethane formation rates

The following cultivable methanogens were tested: *M. acetivorans, M. mazei, M. maripaludis, M. okinawensis* and *M. marburgensis*. For each of those, the standard assay (A) and our optimized assay (B) was employed (**Figure 2)**. For *M. mazei*, the cells were harvested at OD600 value between 1.4-1.6, because a too low protein concentration has been found in lysate prepared from cells harvested at OD600 between 0.4-0.6. For *M. okinawensis*, the cell-free lysates appeared to be almost inactive under both assay conditions.

**Figure 2.**
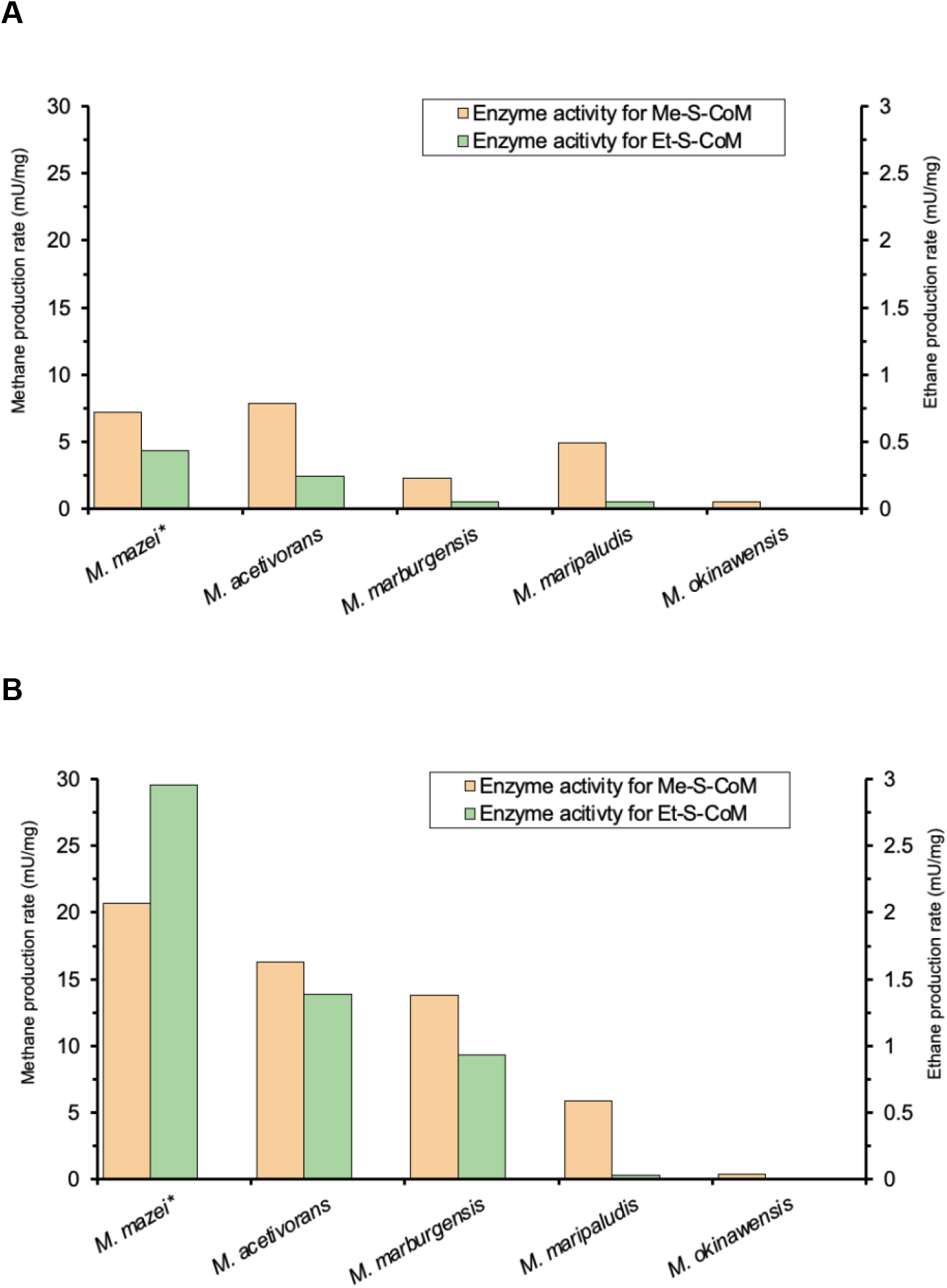
Comparing methane and ethane production rates from different cell extracts in two different assays. Methane from methyl-CoM (left vertical axis) and ethane from: **(A)** Conventional conditions: 37 °C, 0.2 mM B12, 20 mM Ti (III) citrate, and OD600 at harvesting between 0.3-0.5; **(B)** Optimized conditions: 49 °C, 1.1 mM B12, 20 mM Ti(III) citrate, and OD600 at harvesting between 0.3-0.5. *For *M. mazei*, due to low protein concentration in cell lysate, the cells were harvested at OD600 between 1.4-1.6. For enzyme activity, U refers to μmol of (m)ethane formed per minute; mg refers to the total amount of protein in the cell extract.

### Testing the influence of growth conditions on the substrate promiscuity in cell extract from *Methanothermobacter marburgensis*

The high relative ethane-to methane-production rates found for *M. marburgensis* cell-extracts from serum vials (figure 2) prompted us to test cell-extracts from bioreactor grown cells. Cells were grown at 10 L scale with a continuous flow of H_2_/CO_2_ [8,16] until OD_600_ = 5-6. Upon lysis, the cell-free extracts were retrieved and stored at - 80 ºC under H_2_ until further uses. We then conducted assays of Mcr activity using either methyl- or ethyl-S-CoM separately, and in addition we carried out assays with different mixtures of methyl- and ethyl-S-CoM. The conventional assay conditions (0.2 mM cobalamin concentration, 20 mM Ti(III) citrate) and incubation temperature (37 °C) as described by Scheller et al [8,16] were implemented to compare the enzyme activity for methyl-& ethyl-S-CoM (**Figure 3**).

**Figure 3.**
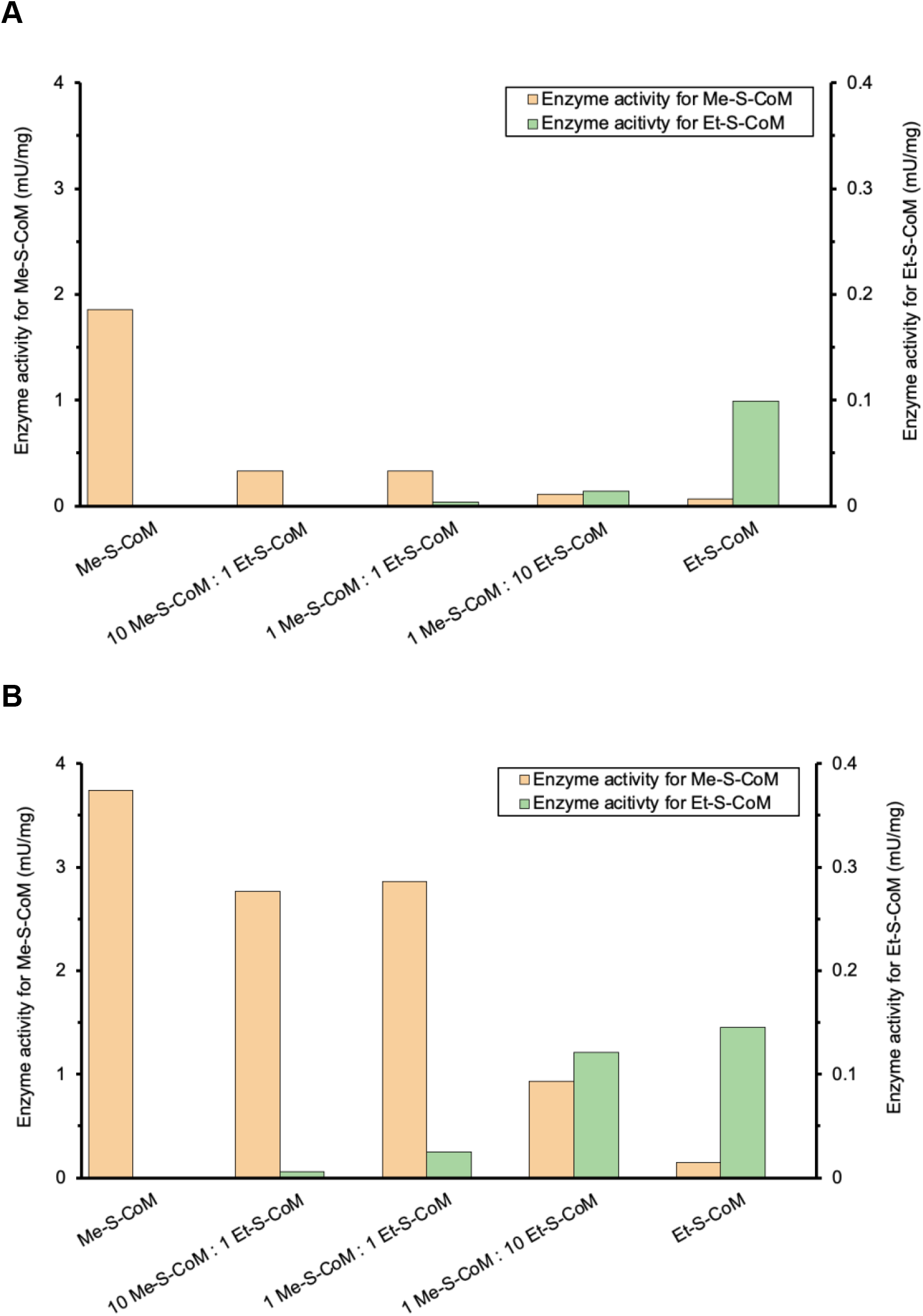
Comparison of enzyme activity for methane (left vertical axis) and ethane production (right vertical axis) from mixtures of methyl-coenzyme M and ethyl-coenzyme M by cell-free lysates from *M. marburgensis* grown under two different conditions. **(A)** Bioreactor-grown samples **(B)** Culture grown in serum vial with a periodic fill of growth substrate under high pressure. The experiments were carried out under standard conditions: 65 °C, 0.2 mM B12, 20 mM Ti (III) citrate, and OD600 at harvesting between 0.3-0.5. For enzyme activity, U is referred to μmol of product formed per minute; mg refers to the total amount of protein in the cell extract.

Cell-free lysate of *M. marburgensis* grown in serum vials to low-OD600 values showed a slightly higher relative enzyme activity for ethane formation. In the experiments where mixtures of methyl- and ethyl-S-CoM have been provided, it appears that ethyl-S-CoM inhibited methane formation for cell-exctracts from the bioreactor (**Figure 3A**), whereas cell-free lysate harvested at low OD showed less inhibition of ethyl-S-CoM on methane formation (**Figure 3B**).

To check whether this difference in enzyme activity depending on growth conditions is related to the differential expression of the two isoenzymes I and II, we partially purified Mcr from each type of cell extracts following the protocol described by Rospert et al. [23]. The two Mcr isoforms can be distinguished basing the size difference of the gamma subunit *via* SDS-PAGE analysis, since their sizes are 38 kDa for isozyme I and 33 kDa for isozyme II [24]. The bioreactor sample grown to high OD showed three main bands corresponding to Mcr enzyme I [23,24] (**Figure S16**). In the case of the culture grown in the vial to low OD, a faint band corresponding to the Mcr II gamma subunit (33 kDa) is visible, which was absent in the bioreactor sample. The ratio of Mcr I to Mcr II can be roughly estimated to be ca. 3:1 for Mcr I to Mcr II based on the intensity of the two gamma subunit bands in the vial sample.

### Discussion

### Utilization of cell-lysates to study the Mcr-catalyzed methane- and ethane formation

We were able to estimate the relative rates of methane formation and of ethane formation from cell extracts of different methanogens. For cell-free lysate of *M. acetivorans* harvested at OD_600_ = 0.3-0.5, we found that including a higher vitamin B_12_ concentration (1.1 mM) and a higher temperature (49 °C) afforded a higher enzyme activity (Assay conditions **B**). For methane formation, the activity increased from 7.5 mU/mg to 16.5 mU/mg, and for ethane formation from 0.25 mU/mg to 1.35 mU/mg. The optimized assay **B** also indicates that Mcr is thermostable at temperatures above the temperature limit for growth of the organisms.

The Mcr activity of *M. acetivorans* cell lysate (7.5 mU/mg) was significantly lower than that reported in a previous study (c.a. 420 mU/mg) [22], which we attribute to the freezing and storing the cell lysates at -80 °C, which has been done in this study in order to facilitate the experiments, allowing to first prepare all cell lysates and then measure the rates. Cell-free lysates *M. maripaludis* showed only a low activity with our assay conditions.

*M. mazei* also showed the highest Mcr activity for methane formation, and the highest substrate promiscuity (ethane formation with 3 mU/mg).

### Mcr activity of cell-free lysates from *M. marburgensis*

Cell-extracts of *M. marburgensis* grown under non-hydrogen limitation, in which Mcr-isoenzyme II is more highly expressed, show a relative ethane-to methane-formation rate of ca. 7.7 % is observed. This rate is much higher than the relative rate of ca. 0.3% reported for purified Mcr-isoenzyme I [15]. Mixtures of methyl- and ethyl-coenzyme M indicate and inhibitory effect of ethyl-coenzyme M for methane formation for cells predominantly expressing Mcr-I, which is in line with the previous finding that ethyl-coenzyme M is readily processed by Mcr, but ethane formation is hindered in this enzyme variant.

## Conclusions

Cell-extracts have been demonstrated to be a suitable tool to estimate the activity of Mcr from various methanogens, without the need for purifying Mcr in its active form. We show that the substrate promiscuity of Mcr towards catalyzing ethane-formation is generally much higher than anticipated, since most reported studies have been carried out with purified Mcr-I from *M. marburgensis*, which appears to be highly selective for methane formation. We have attempted the conversion of propyl-S-CoM towards propane formation, employing cell-free lysates from *M. mazei, M. acetivorans*, and *M. marburgensis* under both assay conditions A and B, but we were not able to detect propane (detection limit = 3 nmol).

## Materials and Methods

### Chemical and Materials

Methyl-S-CoM, ethyl-S-CoM, and coenzyme B were synthesized by following the reported protocol [18]. Gas formation was recorded on an Agilent 6890 GC-FID system (Agilent Technologies, CA, USA) and analyzed using OpenLab CDS software (Agile Technologies, CA, USA). OD600 value and Bradford assay were recorded on BioPhotometer Plus 6132 spectrophotometers (Eppendorf, Germany) at 595 nm wavelength. All chemicals were purchased from Merck. Titanium (III) citrate were prepared following the description elsewhere [25]. MilliQ water was produced by Milli-Q Direct (Merck Millipore, MA, USA). Universal Hood II-Gel Doc system (Bio-Rad) was used to display the SDS-PAGE gel.

### Cultivation of strains

*M. barkeri* (strain Fusaro), *M. acetivorans* (strain WWM73) [21], and *M. mazei* (strain Gö I) [21], *M. maripaludis* (strain JJ:*Δupt*) [26], *M. okinawensis* [27] and *M. marburgensis* [8], were used for this study. For *M. acetivorans*, cell pellets were obtained at different OD600 values. For the other strains, cell pellets were obtained at OD600 value between 0.3-0.5.

For *M. acetivorans*, cultures were first grown statically in High-Salt (HS) medium supplemented with 150 mM MeOH as substrate and 1 mM Na_2_S as sulfur source at 37 °C for 1 day [28]. Typical OD_600_ reaches 0.18-0.20. For low OD_600_ experiments, cultures were then harvested by centrifugation, resuspended in HS medium to yield the cell pellet. For moderate and high OD_600_ experiments, cultures were grown at 25 °C (for moderate OD_600_) or 37 °C (for high OD_600_) until OD_600_ reached the desired value range. The cell pellet was then obtained by centrifugation.

For *M. barkeri*, the desired OD_600_ value (0.3-0.5) can be reached within 3-4 days by statically cultivating at 37 °C in HS medium supplemented with H_2_ and 1% of H_2_S as growth substrate and sulfur source, respectively. For *M. marburgensis*, the cell pellet was obtained using reported cultivation method [28] under H_2_/CO_2_ with H_2_S as growth substrate and sulfur source, respectively. For *M. mazei*, the HS medium was used as growth liquid supplemented with 125 mM MeOH as growth substrate and 1% H_2_S as sulfur source [29]. For *M. maripaludis and M. okinawensis*, the liquid McFc medium was used for cultivation [30,31].

### Cell-free lysate preparation

Cell-free lysate was prepared by sonication (Bandelin Sonopuls, Germany) at 30% amplitude, 30 s pulse and 30 s rest time for 10 min. The cell-lysate fraction was then retrieved by centrifugation and the protein concentration was determined by Bradford assay [31], and expressed in µg/mL. Cell-free lysates were then aliquoted in 10 mL serum vials, of which the headspace was filled with 100% H_2_ and then stored at -80 °C until further uses.

### GC-quantification of methane and ethane

The gas formation was monitored by injecting 100 µL of the head space into the GC-FID. External calibration of methane and ethane formation were conducted for quantification. The method of gas quantification was described elsewhere [22].

### Enzyme activity assay employing cell-free lysate

#### Optimization of enzyme assay

The enzyme assay was performed following previously published protocol [17]. In brief, the assay solution was prepared in 10 mL serum vial. The assay compositions constituted of 2 mL of mixture containing 5 mM alkyl-S-CoM, 0.5 mM coenzyme B homodisulfide (CoB-S-S-CoB), 0.2-1.1 mM of cobalamin, 20 mM of Ti(III) citrate, Tris-HCl (50 mM, pH 7), and 100 µg protein cell-free lysate (prewarmed at RT for 30 min prior to the assay). The solution was then mixed by pipetting and then closed with a rubber stopper. The head space was flushed with 100% H_2_ and then immediately transferred into a heat block at different temperatures (25, 37, and 49 °C) for incubation.

#### Enzyme activity assay of *M. marburgensis* under optimal conditions

For the methyl- and ethyl-S-CoM mixture assay, the previously described protocol was employed with slight modification. The assay solution was prepared in 10 mL serum vial. The assay compositions constituted of 2 mL of mixture containing 0.5 mM Coenzyme B homodisulfide, 0.2 mM of cobalamin, 20 mM of Ti(III) citrate, Tris-HCl (50 mM, pH 7), and 100 µg of cell-free lysate (gradually warmed at RT for 30 min. prior to the assay) with different methyl- and ethyl-S-CoM concentrations were supplemented as following: for only methyl-S-CoM–5mM of methyl-S-CoM; for 9 equivalence (Eq) methyl-S-CoM:Eq ethyl-S-CoM–4.5 mM of methyl-S-CoM and 0.5 mM of ethyl-S-CoM; for 1 Eq methyl-CoM:1 Eq ethyl-S-CoM–2.5 mM of methyl-S-CoM and 2.5 mM of ethyl-S-CoM; for 1 Eq methyl-CoM:9 Eq ethyl-S-CoM–0.5 mM of methyl-S-CoM and 4.5 mM of ethyl-S-CoM; for only ethyl-S-CoM: 5 mM of ethyl-S-CoM. The solution was mixed by pipetting and then closed with a rubber stopper. The headspace was flushed with 100% H_2_ and then immediately transferred into a heat block at 65 °C) for incubation. The gas formation was monitored by injecting 100 µL of the head space into the GC-FID.

### SDS-PAGE analysis of cell-free lysates of *M. marburgensis* harvested under different growth conditions

A fractionated ammonium sulfate precipitation of the *M. marburgensis* lysates was performed as described elsewhere [23]. The cell extracts were first well mixed with ammonium sulfate reaching a 70 % salt concentration and kept on ice for 2 h. After centrifugation (14,000 rpm, 5 min, 4 °C) the supernatants were further enriched with ammonium sulfate until the salt concentration reached about 100 %. After a second centrifugation step the pellets were resuspended in 10 µl sterile water. The protein concentration was estimated by the NanoDrop™ Lite Spectrophotometer (Thermo Fisher). The samples were subsequently mixed with loading dye and incubated for 5 min at 95 °C.

For SDS PAGE a 10 % polyacrylamide gel was loaded with 2 µg protein/lane for both samples. Precision Plus Protein Dual Color Standard (#1610374, Bio-Rad) was applied as a size marker. The electrophoresis was performed at 120 V for 80 min before the gel was processed by Coomassie staining.

## Statistical analysis

The results were expressed as the mean ± standard deviation. Replication was conducted in duplication. The significance of each coefficient was determined by using the Student *t-test* and p-value. The ANOVA analysis was conducted at 95% significance level. The analysis software used for this study was Minitab® 21 (version 3.1) (Minitab Inc, State College, Palo Alto, CA, USA.).

## Acknowledgement

The authors would like to acknowledge Norman Adlung (VTT Finland) for guidance in the early development of the project. The authors would like to acknowledge the Academy of Finland (#13326020) and NovoNordisk Foundation (NNF310279) for fundings.

## Author contribution

T.N.: conceptualization and data curation, T.N. & S.K: conducting experiments, T.N., S.K., & S.S.: drafting the manuscript, S.S.: supervising, project administration and funding acquisition.

