## Supplementary Information for "Methane and ethane production rates by methyl-coenzyme M reductase in cell extracts from different methanogens"

Silvan Scheller 0000-0002-0667-9224

V. P. Thinh Nguyen 0000-0002-2499-2900

Sebastian Keller 0000-0001-7477-1201

### Determination of parameters having effects onto Mcr activities

It appeared that selected parameters such as the growth phase at harvesting, the incubation temperature and the $B_{12}$ concentration involved in the CoB-S-S-CoM heterodisulfide reduction have significant effects on Mcr enzyme activity for methyl-CoM. In order to study these effects, we devised a response surface design involving 3 factorial variables ($OD_{600}$ at harvesting, incubation temperature and cobalamin concentration) that included, not only the enzyme activity for methyl-CoM as a response variable, but also comprised the enzyme activity for ethyl-CoM, and the promiscuity (ratio between ethane and methane formed from ethyl-CoM and methyl-CoM, respectively) to assay the ethane formation ability of studying different methanogens.

#### Growth phase

Cell lysates of different growth phases, with OD600 at harvesting were categorized as early (low OD, value between 0.1-0.3), middle (moderate OD, value between 0.5-0.7) and late growth phase (high OD, value between 0.9-1.2). Mcr assays employing these cell lysates were conducted to determine the effects of growth phase on methanogenic activities. Methyl-CoM was used as an alkyl donor, and methane was quantified by GC-FID using methane as an external standard curve. Our results indicated that *M. acetivorans* Mcr was most active at mid growth phase (OD600 value between 0.5-0.8).

#### Incubation temperature and cobalamin concentration

We investigated the effects of incubation temperature during the assay onto the enzyme activity by conducting Mcr assay at different temperatures (25 and 37 °C) employing different cell-free lysates prepared at different growth phases (low and high OD levels). Our results presented in **Figure 2** shows that low OD culture exhibited higher enzyme activity at 37 °C.

Ti(III) citrate is added with cyanocobalamin to reduce the generated CoB-S-S-CoM heterodisulfide complex in the Mcr activity assay [12,13]. The concentration of Ti(III) must be in excess, whereas the cobalamin concentration was reported to be between 0.2-2 mM to ensure the highest Mcr activity [12]. Taken together, we hypothesized that cobalamin concentration might have an indirect effect on Mcr assay. Therefore, the concentration of cobalamin–the source of (cyano)cobalamin in our assay–was also included as a factorial variable for our design of experiment. We investigated the concentration range of cobalamin between 0.2 to 1.1 mM, as higher concentration might not be directly beneficial to our assay. Our results depicted in **Figure 2** showed that low OD and low cobalamin concentration had significant positive effect onto enzyme activity for methyl-CoM, whereas increasing cobalamin concentration only had a slight declination of the enzyme activity for high OD cell-free lysate. These results confirmed our hypothesis concerning the indirect effect of cobalamin onto Mcr enzyme activity.

Based on our previous results, we proceeded the independent optimization of the Mcr enzyme activity with a non-C1 substrate, ethyl-CoM, to determine the enzyme activity of Mcr in cell-free lysates towards more extended substrate (C2 in this study), as well as the promiscuity of Mcr towards these substrates. For the latter purpose, we determined the ratio between ethane and methane formed by separately providing methyl-CoM and ethyl-CoM as alkyl donors were therefore determined. Response surface methodology is a powerful statistical technique that has been successfully employed to optimize the methanogenic activity of methanogens [22]. Thereby, we believed that employing this statistical approach would afford an efficient and highly reproducible procedure that enables alkane formation from corresponding alkyl-CoM. The outcomes of this optimization process would allow us to investigate the enzyme activity for ethyl-CoM and promiscuity of cell-free lysate from *M. acetivorans*. The model fitting and further optimization of enzyme activity are presented in **Supporting Information**. Confirmatory experiments were conducted in duplicate according to the operating conditions afforded *via* the prediction model to confirm the estimating values. Although the experimental values were higher than the predicted value, these remained satisfactory in the context of our study.

### Optimization of Mcr assay employing Box-Benken Response Surface design

Interaction between different parameters onto enzyme activity and promiscuity

Based on the outcomes of the investigation of parameters having effects onto Mcr activities, we proceeded with the optimization of the Mcr enzyme activity with a non-C1 substrate, ethyl-CoM, in parallel in order to determine the enzyme activity of Mcr in cell-free lysates towards more extended substrate ($C_2$ in this study), as well as the promiscuity of Mcr towards these substrates. For the latter purpose, we determined the ratio between ethane and methane formed by separately providing methyl-CoM and ethyl-CoM as alkyl donor was therefore determined. Response surface methodology is a powerful statistical technique that has been successfully employed to optimize the methanogenic activity of methanogens [1]. Thereby, we believed that employing this statistical approach would afford an efficient and highly reproducible procedure that enables alkane formation from corresponding alkyl-CoM. The outcomes of this optimization process would allow us to investigate the enzyme activity for ethyl-CoM and promiscuity of cell-free lysate from *M. acetivorans*. A 3-factor design was therefore generated for this purpose (**Table S1**). The model fitting and further optimization of enzyme activity are presented in the following subsection.

A Box-Behnken design was used to optimize the assay conditions of Mcr activity using methyl- and ethyl-CoM as substrates, with a total of 13 experiments and a triplicate at the central point (**Table S1**). Enzyme activity for methyl- ($Y_1$), ethyl-CoM ($Y_2$) and the ratio between ethane and methane formation from methyl- and ethyl-CoM (referred as the promiscuity, $Y_3$) were determined as the responses, whereas the independent variables used in this optimization at three different levels (-1, 0, +1) were defined and studied as following: $OD_{600}$ at harvesting ($X_1$), incubation temperature ($X_2$), and vitamin $B_{12}$ concentration ($X_3$). **Table S5** described different levels of selected factors for the optimization process.

Calculation methods of selected responses are described in **Table S6**. For the calculation of $Y_1$ and $Y_2$, gas formation at 4 timepoints (90, 180, 240 and 300 mins) were consistently monitored and the outcome values were used for the construction of linear regression of the enzyme activity.

The experimental data were recorded and then fitted using a second-order polynomial equation:

$$Y_q = \beta_0 + \sum_{i=1}^{2} \beta_i X_i + \sum_{i=1}^{2} \beta_{ii} X_i^2 + \sum_{i<j=1}^{2} \beta_{ij} X_i X_j + \varepsilon_{residues\ q'} \tag{1}$$

$Y_q$ is the different responses (q = 1-3), $\beta_0$, $\beta_i$, $\beta_{ii}$ and $\beta_{ij}$ are the regression coefficients of the mean, linear, quadratic and interaction terms, respectively; $X_i$ and $X_j$ are the independent variables; and $\varepsilon_{residues\ q'}$ is the difference between the observed and the predicted value.

Model Fitting

Multiple regression equations using polynomial second order were applied to afford the responses. The regression coefficients of each model were analyzed by ANalysis Of Variance (ANOVA) and reported in **Table S2-4**. In overall, each generated models were validated by ANOVA (model p-value < 0.05) accompanied with the lack-of-fitted test (p-value > 0.05), with non-significant factors were removed for model refitting purposes. The accuracy of our models was furthermore confirmed according to the $R_2$ and $R_2$-adjusted value comparing to $R_2$ predicted values.

#### Optimization of enzyme activity and promiscuity towards C2-substrate

According to our results depicted in **Figure S1**, all of selected factors including OD at harvesting, incubation temperature and $B_{12}$-concentration in the activity assay have significant effects onto the conversion of methyl-CoM towards methane, and the promiscuity of cell-free lysates from *M. acetivorans*. In contrast, only OD at harvesting and incubation temperature are significant factors for the enzyme activity for ethyl-CoM towards ethane formation. Moreover, the significant interaction and quadratic terms between OD and incubation temperature were observed according to generated models. The effects of OD and incubation temperature on enzyme activity towards ethyl-CoM and promiscuity (**Table S3** and **Table S4**) suggest that the optimal OD600 at harvesting should be at the moderate value (between 0.4-0.8) in order to maintain Mcr enzyme activity. Nevertheless, the model of promiscuity also showed that low OD resulted in better ratio between ethane and methane formation from ethyl- and methyl-CoM, respectively. Thereby, a compromise between the enzyme activity for ethyl-CoM and the promiscuity must be reached in order to implement this assay method to screen for the promiscuity of other methanogens. In this context, an optimization employing analytical procedure was conducted to afford the optimal operating conditions for this assay.

Our results of optimization procedure (**Figure S2**) suggested that the optimal assay conditions to maximize the promiscuity and enzyme activity towards ethyl-CoM and to minimize the enzyme activity towards methyl-CoM is as following: Moderate/low $OD_{600}$ at harvesting (0.4-0.5), high incubation temperature (49 °C) and high $B_{12}$ concentration (1.1 mM) (**Figure S5**). Under these conditions, the enzyme activity towards methyl-CoM, ethyl-CoM and the promiscuity are predicted to be 8.4 mU/mg, 1.4 mU/mg and 10.9%, respectively, with U is referred to µmol of product formed per minute.

**Table S1.** Results for Box-Behnken Design for 15 samples of Mcr enzyme assay

| Entry | Run order | Factors | | | Responses | | |
|---|---|---|---|---|---|---|---|
| | | OD600 $X_1$ | Incubation temperature (°C) $X_2$ | Vitamin B12 concentration (mM) $X_3$ | Methane (U/mg) $Y_1$ | Ethane formation (U/mg) $Y_2$ | Ratio Ethane/Methane (%) $Y_3$ |
| 1 | 1 | Medium | 49 | 1,1 | 0,00966 | 0,00116 | 11,99793 |
| 2 | 3 | Low | 25 | 0,65 | 0,00078 | 0,00002 | 2,67738 |
| 3 | 8 | High | 25 | 0,65 | 0,00059 | 0,00000 | 0,20769 |
| 4 | 4 | Low | 49 | 0,65 | 0,00519 | 0,00027 | 5,14368 |
| 5 | 9 | High | 49 | 0,65 | 0,00111 | 0,00002 | 1,60342 |
| 6 | 11 | Low | 37 | 0,2 | 0,00163 | 0,00017 | 10,23385 |
| 7 | 10 | High | 37 | 0,2 | 0,00058 | 0,00005 | 8,78402 |
| 8 | 12 | Low | 37 | 1,1 | 0,00132 | 0,00011 | 8,18665 |
| 9 | 13 | High | 37 | 1,1 | 0,00126 | 0,00001 | 0,73217 |
| 10 | 5 | Medium | 25 | 0,2 | 0,00095 | 0,00007 | 4,21335 |
| 11 | 6 | Medium | 49 | 0,2 | 0,01870 | 0,00125 | 6,68449 |
| 12 | 7 | Medium | 37 | 0,65 | 0,00579 | 0,00031 | 5,38687 |
| 13 | 14 | Medium | 37 | 0,65 | 0,06870 | 0,00168 | 2,44833 |
| 14 | 2 | Medium | 25 | 1,1 | 0,05270 | 0,00004 | 0,08059 |
| 15 | 15 | Medium | 37 | 0,65 | 0,00525 | 0,00032 | 6,05524 |

**Table S2**. Analysis of variance for the fitted regression model of methane formation (Y1).

| Analysis of Variance for Transformed Response | | | | | |
|---|---|---|---|---|---|
| **Source** | **DF** | **Adj SS** | **Adj MS** | **F-Value** | **P-Value** |
| Model | 5 | 2172,96 | 434,59 | 17,17 | <0.001 |
| **Linear** | 3 | 693,44 | 231,15 | 9,13 | 0,004 |
| OD | 1 | 188,26 | 188,26 | 7,44 | 0,023 |
| Incubation_Temp | 1 | 344,85 | 344,85 | 13,63 | 0,004 |
| B12_Concentration | 1 | 160,34 | 160,34 | 6,34 | 0,032 |
| **Square** | 1 | 1240,44 | 1240,44 | 49,02 | <0.001 |
| OD*OD | 1 | 1240,44 | 1240,44 | 49,02 | <0.001 |
| **2-Way Interaction** | 1 | 239,08 | 239,08 | 9,45 | 0,013 |
| Incubation_Temp*B12_Concentration | 1 | 239,08 | 239,08 | 9,45 | 0,013 |
| **Error** | 9 | 227,76 | 25,31 | | |
| Lack-of-Fit | 7 | 165,38 | 23,63 | 0,76 | 0,674 |
| Pure Error | 2 | 62,38 | 31,19 | | |
| **Total** | 14 | 2400,72 | | | |

| Model Summary for Transformed Response | | | |
|---|---|---|---|
| **S** | **R-sq** | **R-sq(adj)** | **R-sq(pred)** |
| 0,843702 | 90,51% | 85,24% | 71,99% |

The model predicting the methane formation with unscaled coefficient is shown in **Equation 1**:

$$-Y_1^{-0.5} = -73.3 - 4.85X_1 + 1.478X_2 + 62.3X_3 - 18.23X_1^2 - 1.432X_2X_3 \qquad \text{(Eq 1)}$$

**Table S3.** Analysis of variance for the fitted regression model of Ethane formation (Y2).

| **Analysis of Variance for Transformed Response** | | | | | |
|---|---|---|---|---|---|
| **Source** | **DF** | **Adj SS** | **Adj MS** | **F-Value** | **P-Value** |
| Model | 3 | 48,872 | 16,2906 | 22,89 | 0 |
| **Linear** | 2 | 28,474 | 14,2369 | 20 | 0 |
| OD | 1 | 10,54 | 10,5396 | 14,81 | 0,003 |
| Incubation_Temp | 1 | 17,934 | 17,9342 | 25,19 | 0 |
| **Square** | 1 | 20,398 | 20,398 | 28,66 | 0 |
| OD*OD | 1 | 20,398 | 20,398 | 28,66 | 0 |
| **Error** | 11 | 7,83 | 0,7118 | | |
| Lack-of-Fit | 9 | 5,958 | 0,662 | 0,71 | 0,708 |
| Pure Error | 2 | 1,872 | 0,9359 | | |
| **Total** | 14 | 56,702 | | | |
| **Model Summary for Transformed Response** | | | | | |
| **S** | **R-sq** | **R-sq(adj)** | | **R-sq(pred)** | |
| 0,843702 | 86,19 % | 82,42 % | | 75,23 % | |

The model predicting the methane formation with unscaled coefficient is shown in **Equation 2**:

$$ln(Y_2) = -12.638 - 1.148X_1 + 0.1248X_2 - 2.337X_1^2 \quad \text{(Eq. 2)}$$

**Table S4.** Analysis of variance for the fitted regression model of promiscuity (Ethane/Methane)

| **Analysis of Variance for Transformed Response** | | | | | |
|---|---|---|---|---|---|
| **Source** | **DF** | **Adj SS** | **Adj MS** | **F-Value** | **P-Value** |
| Model | 5 | 26,2125 | 5,2425 | 12,27 | 0,001 |
| **Linear** | 3 | 17,8905 | 5,9635 | 13,96 | 0,001 |
| OD | 1 | 4,9442 | 4,9442 | 11,57 | 0,008 |
| Incubation_Temp | 1 | 8,3261 | 8,3261 | 19,49 | 0,002 |
| B12_Concentration | 1 | 4,6202 | 4,6202 | 10,81 | 0,009 |
| **Square** | 1 | 3,1655 | 3,1655 | 7,41 | 0,024 |
| Incubation_Temp*Incubation_Temp | 1 | 3,1655 | 3,1655 | 7,41 | 0,024 |
| **2-Way Interaction** | 1 | 5,1565 | 5,1565 | 12,07 | 0,007 |
| Incubation_Temp*B12_Concentration | 1 | 5,1565 | 5,1565 | 12,07 | 0,007 |
| **Error** | 9 | 3,8452 | 0,4272 | | |
| Lack-of-Fit | 7 | 3,36 | 0,48 | 1,98 | 0,376 |
| Pure Error | 2 | 0,4852 | 0,2426 | | |
| **Total** | 14 | 30,0577 | | | |

| **Model Summary for Transformed Response** | | | |
|---|---|---|---|
| **S** | **R-sq** | **R-sq(adj)** | **R-sq(pred)** |
| 0,653636 | 87,21 % | 80,10 % | 57,85 % |

The model predicting the methane formation with unscaled coefficient is shown in **Equation 3**

$$ln(Y_3) = -4.22 - 0.786X_1 + 0.422X_2 - 9.47X_3 - 0.00639X_2^2 + 0.2103X_2X_3 \quad \text{(Eq. 3)}$$

**Table S5.** Factorial variables and levels in the response surface design.

| **Factors** | **Symbol** | **Level** | | |
|---|---|---|---|---|
| | | **-1** | **0** | **+1** |
| OD600 at harvesting | A | 0.1-0.3 | 0.5-0.8 | 1.0-1.2 |
| Incubation temperature | B | 25 °C | 37 °C | 49 °C |
| Vitamin $B_{12}$ concentration | C | 0.2 mM | 0.65 mM | 1.1 mM |

**Table S6.** Symbol and calculation method responses for the response surface design.

| Responses | Symbols | Calculation method |
|---|---|---|
| Enzyme activity towards methyl-CoM (U/mg of protein) | $Y_1$ | Slope of methane formation curve |
| Enzyme activity towards ethyl-CoM (U/mg of protein) | $Y_2$ | Slope of ethane formation curve |
| The promiscuity | $Y_3$ | $Y_1/Y_2 \times 100\%$ |

**Table S7.** Enzyme activity for methyl- and ethyl-CoM, and the promiscuity of cell-free lysate under optimal conditions generated by the prediction models.

| | Activity methyl-CoM (mU/mg) | Activity ethyl-CoM (mU/mg) | Promiscuity (%) |
|---|---|---|---|
| **Prediction value** | 8.4 | 1.5 | 10.9 |
| **Replicate 1** | 16.3 | 1.37 | 8.5 |
| **Replicate 2** | 16.7 | 1.43 | 8.9 |
| **Mean ± S.D.** | 16.5 ± 0.21 | 1.43 ± 0.06 | 8.7 ± 0.2 |

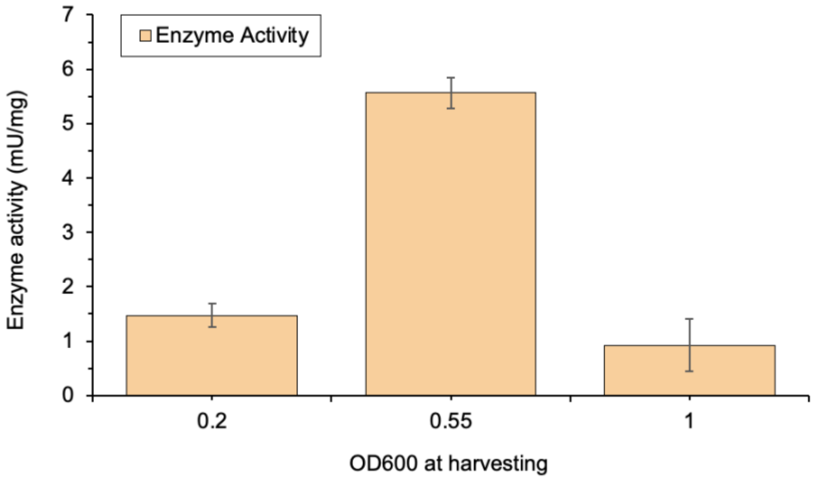


**Figure S1.** Influence of the growth phase onto Mcr methane formation activity from methyl-CoM and Coenzyme B in cell-free lysates prepared from M. acetivorans supplemented with MeOH.

**A**

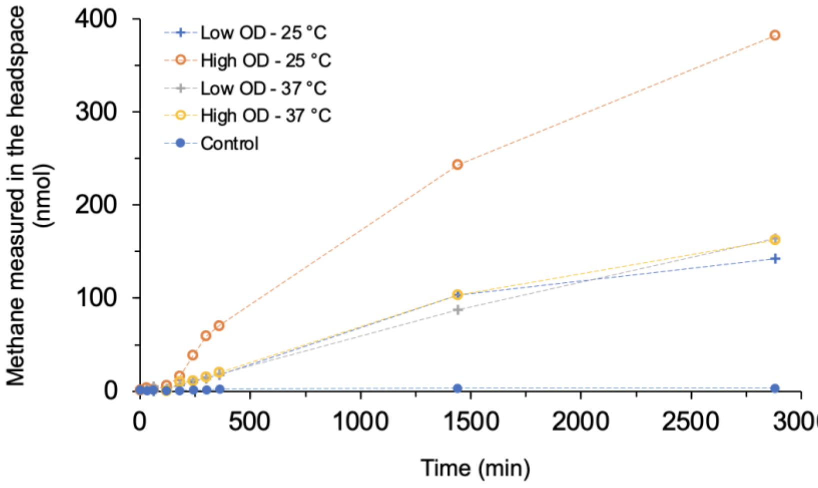


**B**

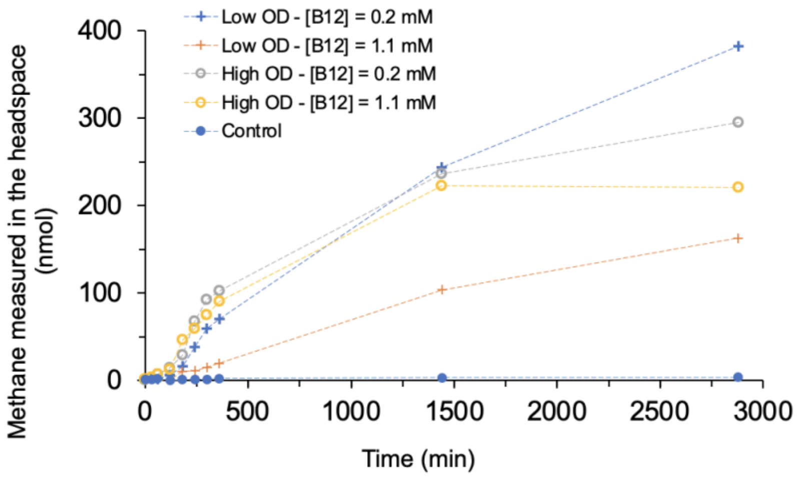


**Figure S2** Methane production of *M. acetivorans* cell-free lysates under different conditions. (A) Varying the incubation temperature (25 and 37 °C) for cells harvested at two different growth phases (low and high $OD_{600}$). (B) Varying the amound vitamin $B_{12}$ added (0.2 mM, 1.1 mM) in experiments ad two different growth phase (low and high $OD_{600}$). Assays were conducted in duplicate under conventional assay conditions (100 µg of protein in cell lysate, 0.2 mM vitamin B12, 0.2 mM $B_{12}$, 37 °C).

**A)** **B)**

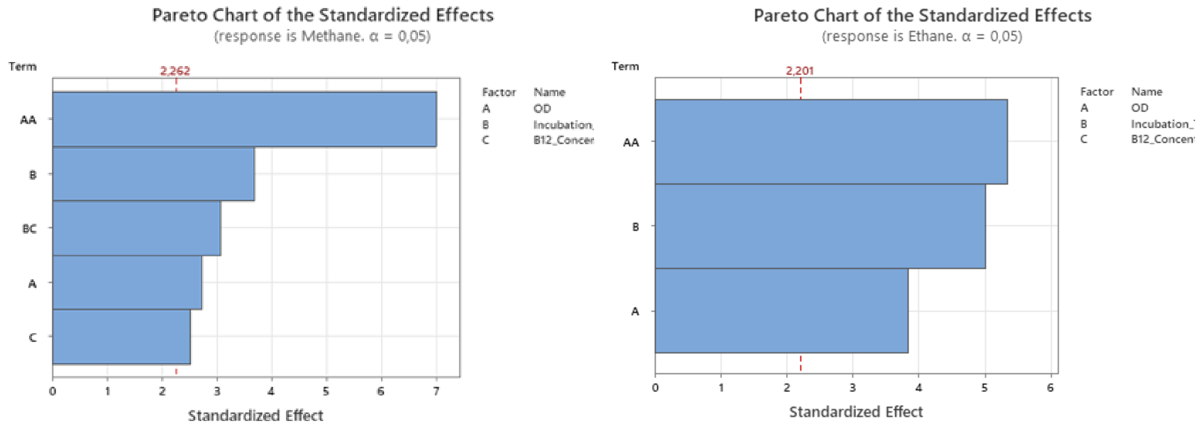


C)

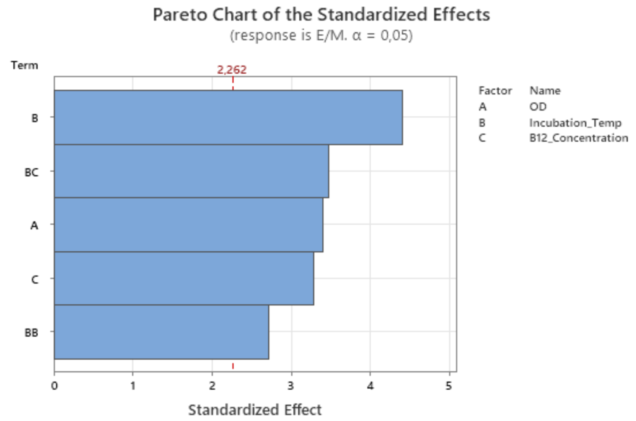


**Figure S3.** Parento charts of the standardized effects for selected responses: A) enzyme activity for methyl-CoM; B) enzyme activity for ethyl-CoM; and C) the promiscuity

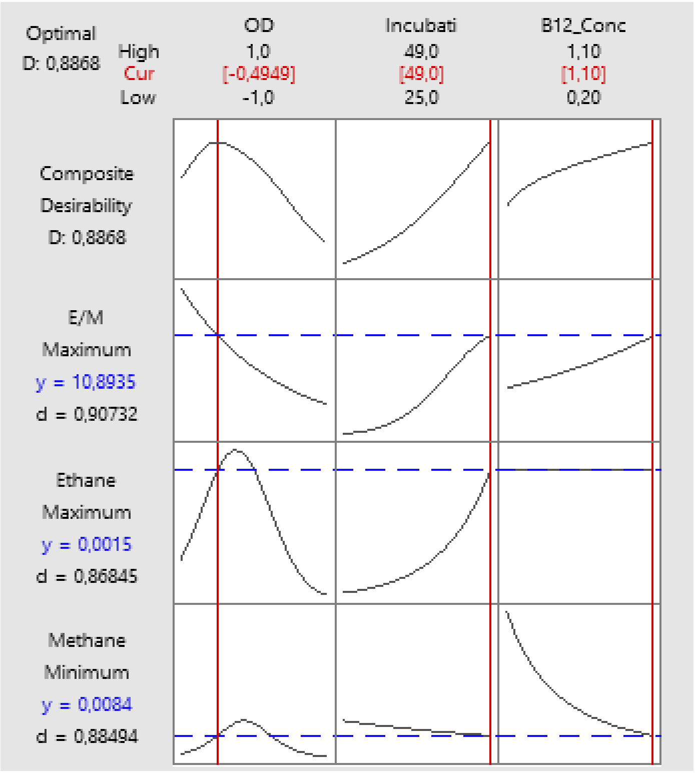

**Figure S4.** Optimization chart. The red traits indicate the optimal conditions for each factor according to our prediction model.

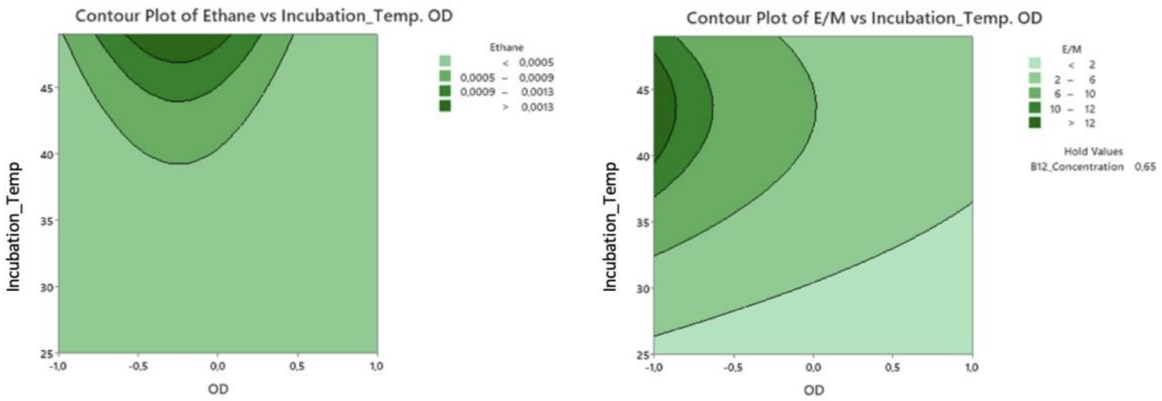


**Figure S5.** Effect of OD600 at harvesting and incubation temperature on enzyme activity for ethyl-CoM and the promiscuity of cell-free lysate of *M. acetivorans* grown on MeOH. Enzyme activity and the promiscuity were respectively expressed in U/mg of protein in cell-free lysate and %. For the contour plot of the promiscuity, the value of $B_{12}$-concentration was determined to be significant and, thereby, was held at 0.65. the promiscuity was expressed in %.

**A)**

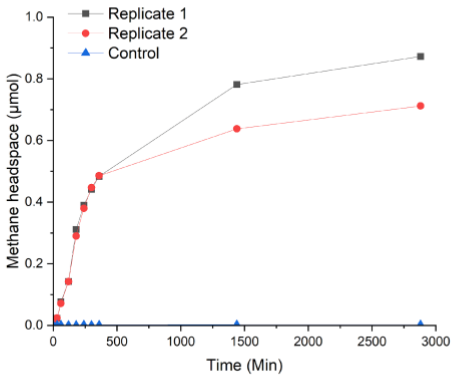


**B)**

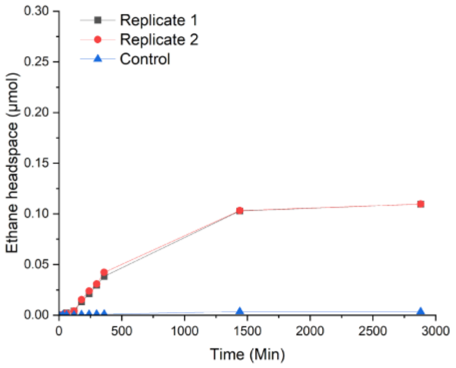


**C)**

**D)**

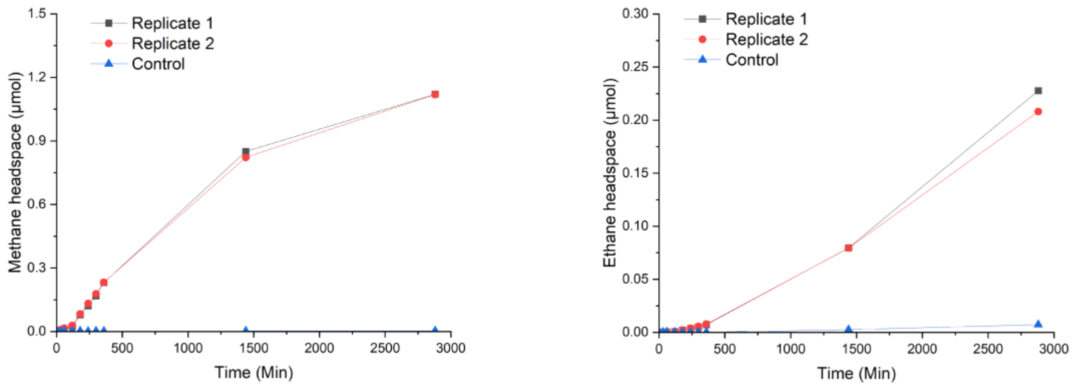


**Figure S6.** Results of confirmatory Mcr assay experiments under optimal (**A** & **B**) comparing to that of conventional operating conditions (**C** & **D**). In A & C, only methyl-CoM (5 mM) was supplemented, whereas ethyl-CoM (5 mM) was sole alkyl-CoM source in **C** & **D**.

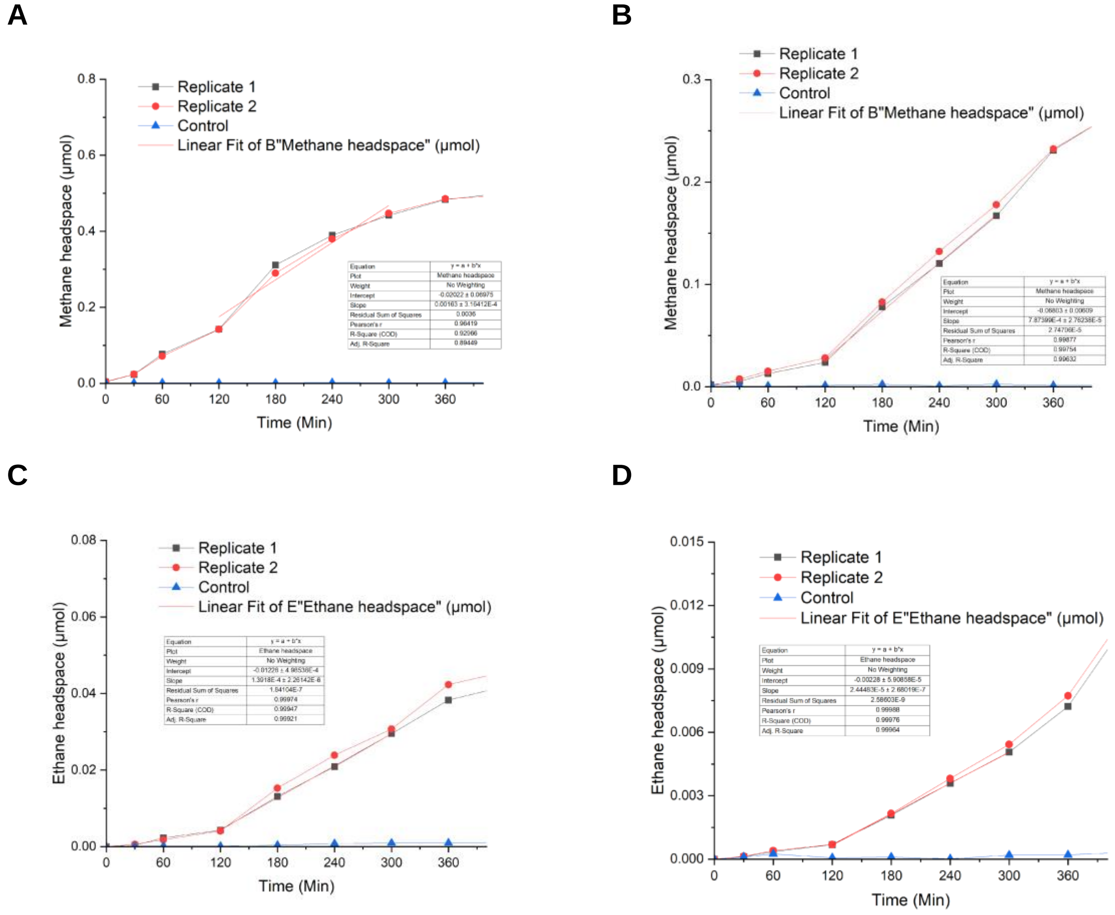


**Figure S7.** Linear regression of gas formation from alkyl-CoM mediated by Methanosarcina. acetivorans cell-free lysate under optimal and conventional conditions. **A** & **B** show the methane formation from methyl-CoM under optimal and conventional conditions, respectively. **C** & **D** show the ethane formation from ethyl-CoM under optimal and conventional conditions, respectively. Four time points (120, 180, 240, 300 mins) were consistently used to determine the enzyme activity for both methyl- and ethyl-CoM.

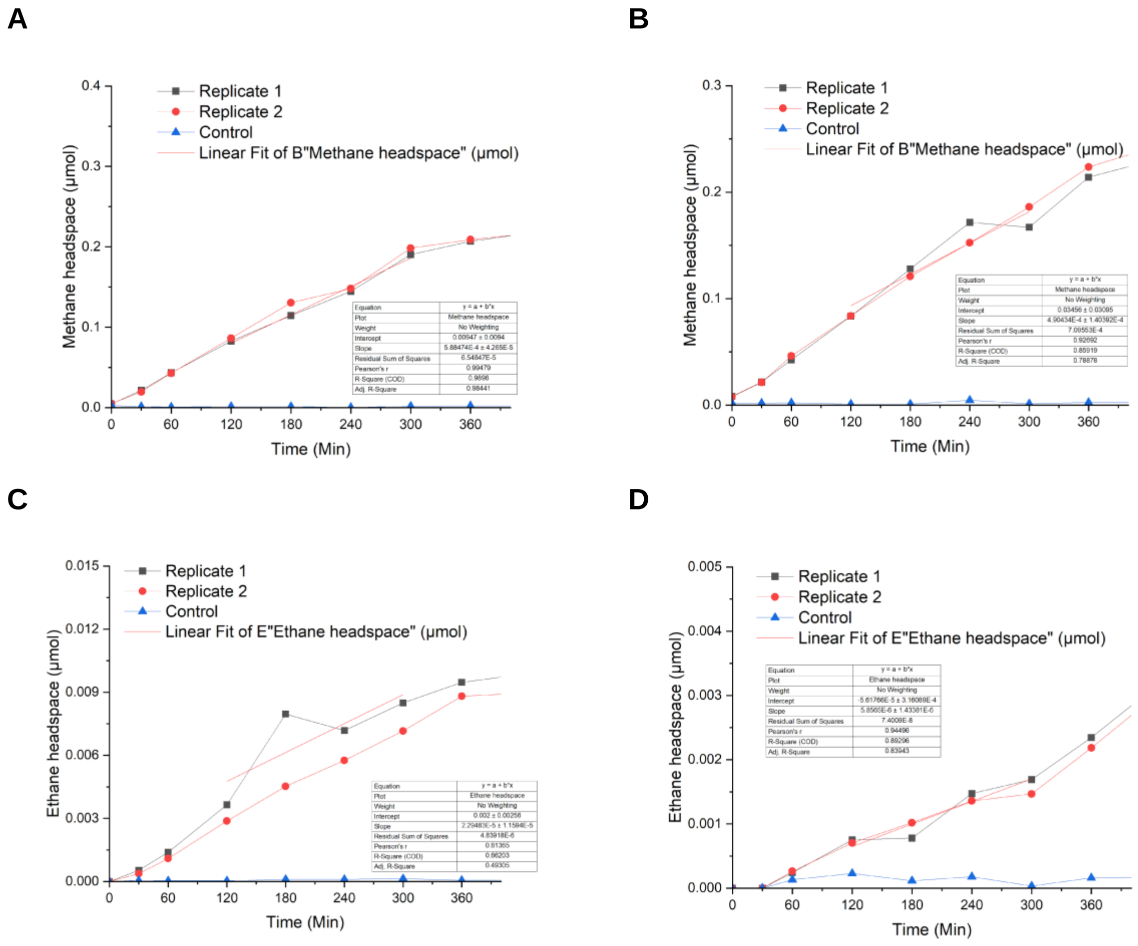


**Figure S8.** Linear regression of gas formation from alkyl-CoM mediated by Methanococcus. maripaludis cell-free lysate under optimal and conventional conditions. **A** & **B** show the methane formation from methyl-CoM under optimal and conventional conditions, respectively. **C** & **D** show the ethane formation from ethyl-CoM under optimal and conventional conditions, respectively. Four time points (120, 180, 240, 300 mins) were consistently used to determine the enzyme activity for both methyl- and ethyl-CoM.

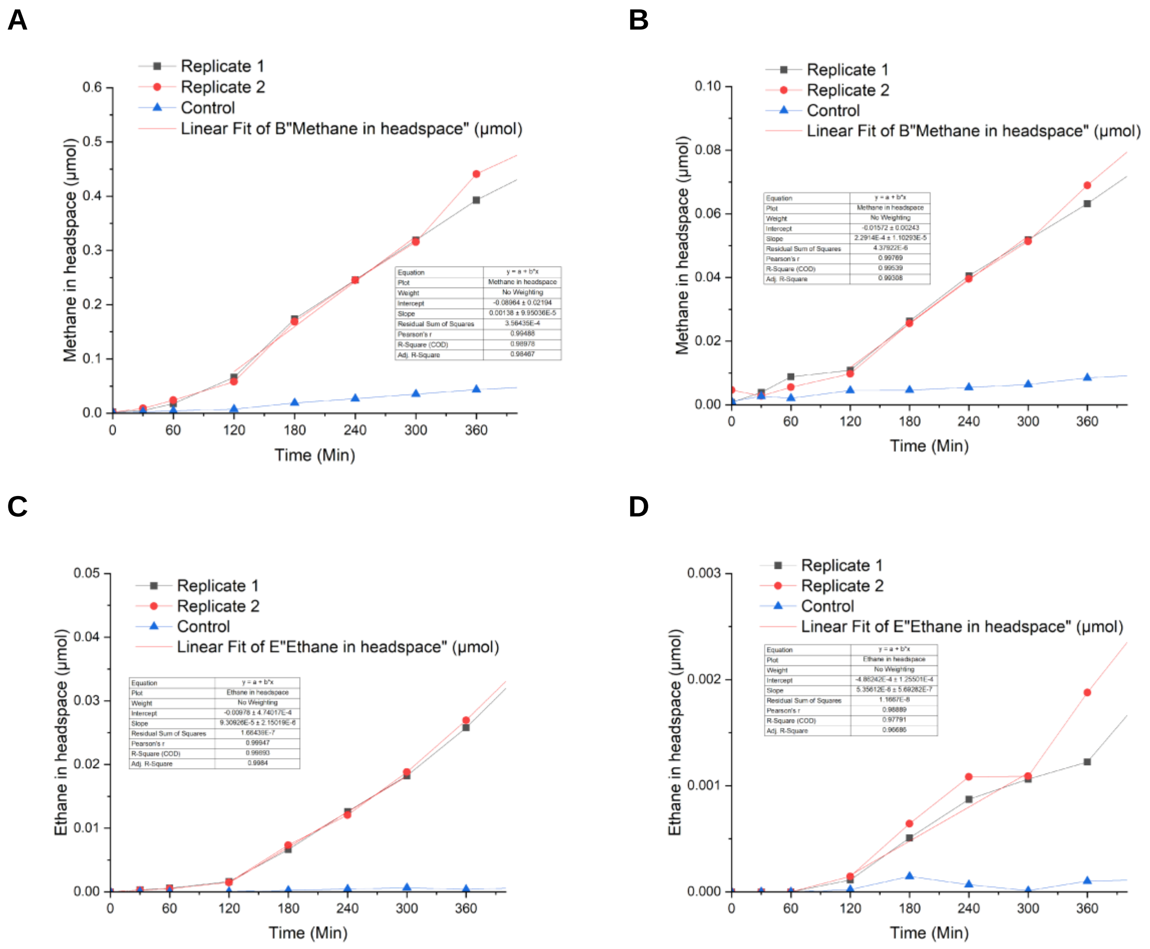


**Figure S9**. Linear regression of gas formation from alkyl-CoM mediated by *Methanothermobacter marburgensis* cell-free lysate under optimal and conventional conditions. **A** & **B** show the methane formation from methyl-CoM under optimal and conventional conditions, respectively. **C** & **D** show the ethane formation from ethyl-CoM under optimal and conventional conditions, respectively. Four time points (120, 180, 240, 300 mins) were consistently used to determine the enzyme activity for both methyl- and ethyl-CoM.

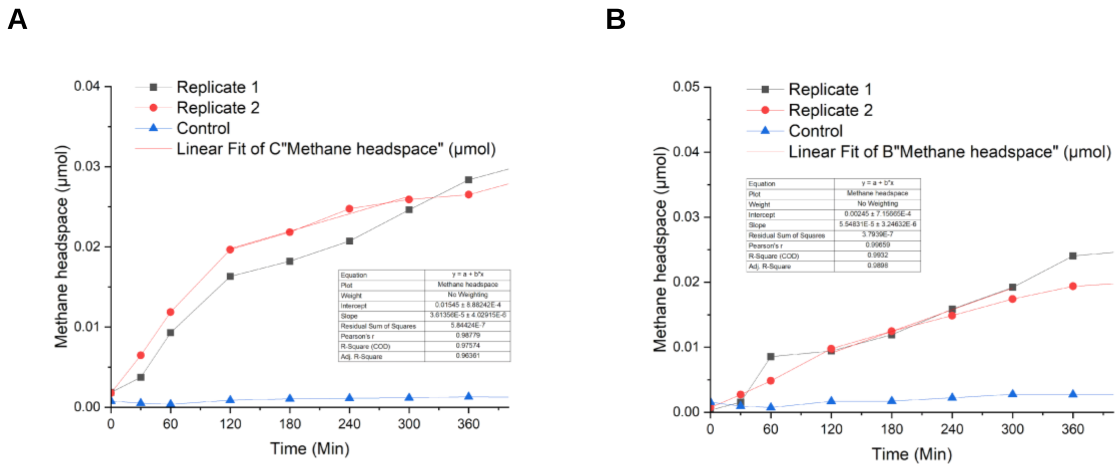


**Figure S10.** Linear regression of gas formation from alkyl-CoM mediated by *Methanothermococcus okinawensis* cell-free lysate under optimal and conventional conditions. **A** & **B** show the methane formation from methyl-CoM under optimal and conventional conditions, respectively. Four time points (120, 180, 240, 300 mins) were consistently used to determine the enzyme activity for both methyl- and ethyl-CoM. No ethane formation from ethyl-CoM using *M. okinawensis* cell-free lysate has occurred.

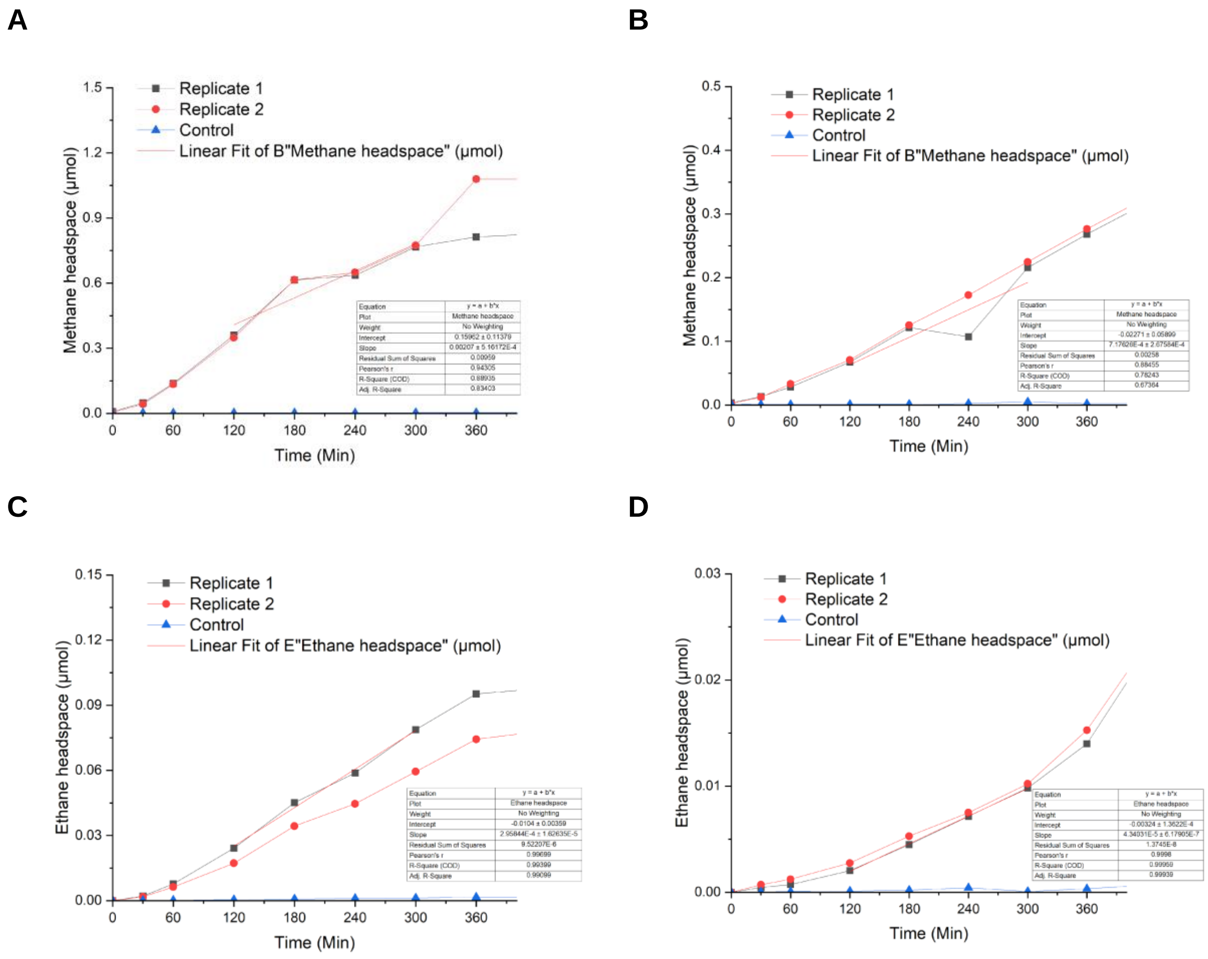


**Figure S11.** Linear regression of gas formation from alkyl-CoM mediated by Methanosarcina mazei grown on MeOH cell-free lysate under optimal and conventional conditions. **A** & **B** show the methane formation from methyl-CoM under optimal and conventional conditions, respectively. **C** & **D** show the ethane formation from ethyl-CoM under optimal and conventional conditions, respectively. Four time points (120, 180, 240, 300 mins) were consistently used to determine the enzyme activity for both methyl- and ethyl-CoM.

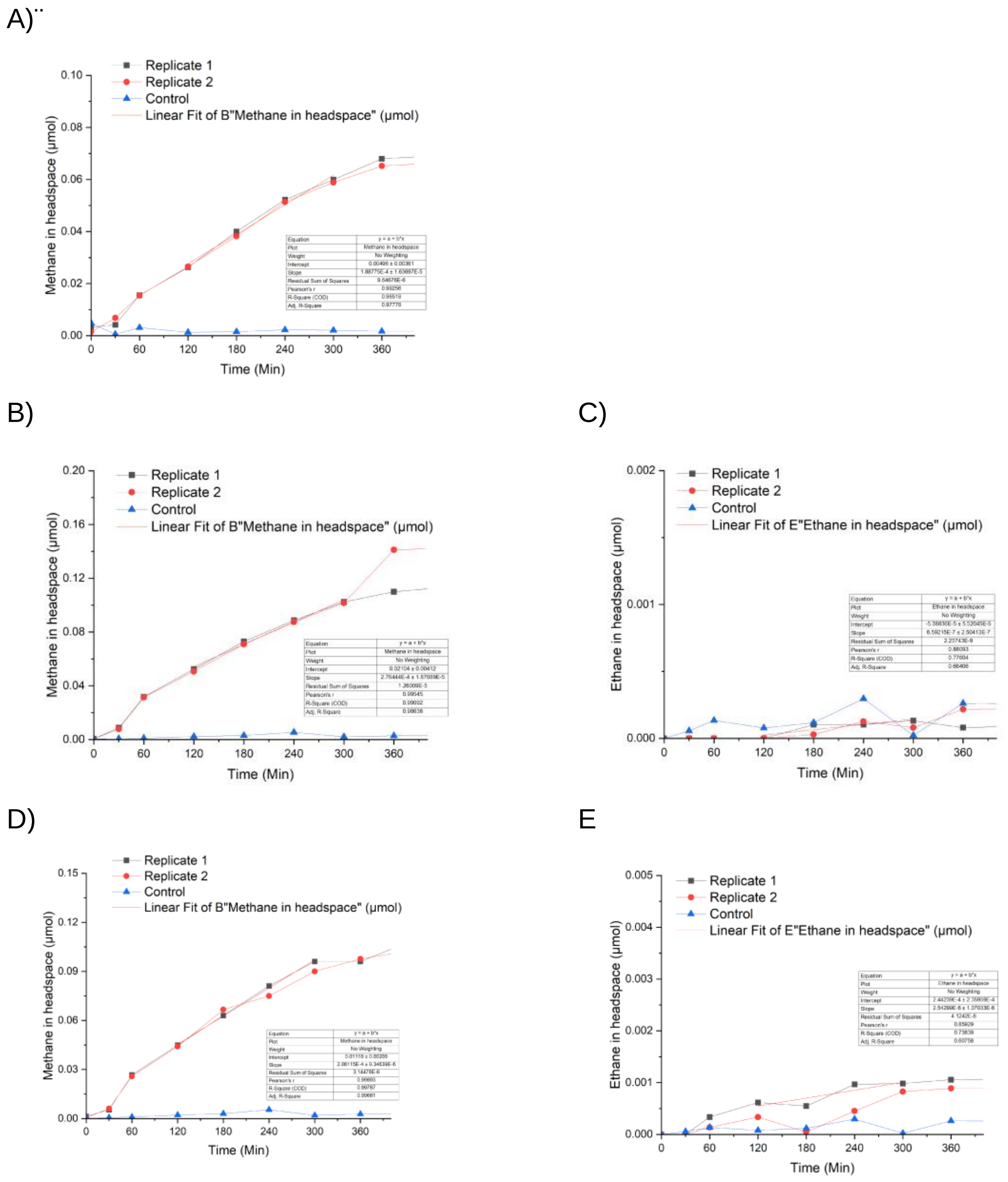


**Figure S12.** Linear regression of gas formation from alkyl-CoM mediated by Methanothermobacter marburgensis grown on $H_2/CO_2$ in serum vial. Mcr assay was conducted at 65 °C using 1.1 mM vitamin $B_{12}$, 20 mM Ti(III) citrate, 0.5 mM of heterodisulfide of coenzyme B, and different concentration of alkyl-CoM as following: **(A)** 5 mM of methyl-CoM; **(B) & (C)** 4.5 mM of methyl-CoM & 0.5 mM of ethyl-CoM; **(D)** & **(E)** 2.5 mM of methyl-CoM & 2.5 mM of ethyl-CoM; **(F)** & **(G)** 0.5 mM of methyl-CoM & 4.5 mM of ethyl-CoM; **(H)** & **(I)** 5 mM of ethyl-CoM. All of the experiments were conducted with 100 µg of cell-free lysate from Mt. marburgensis, except **(A)** where 50 µg of cell-free lysate was used instead.

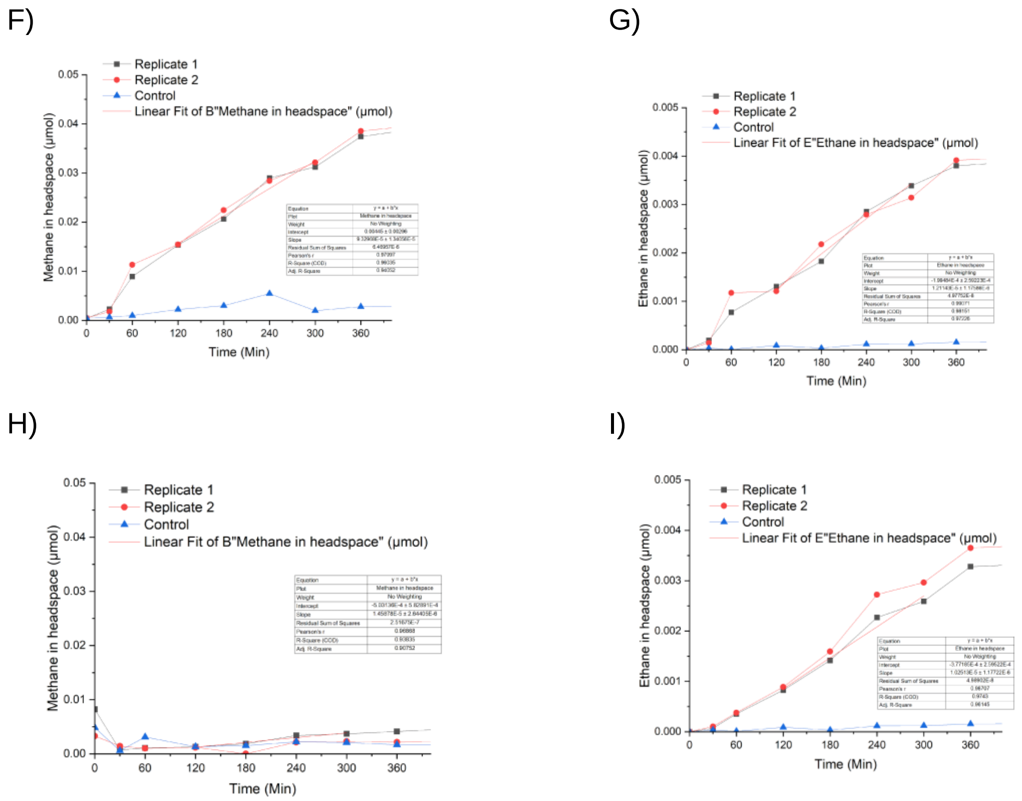


**Figure S12.** (continued).

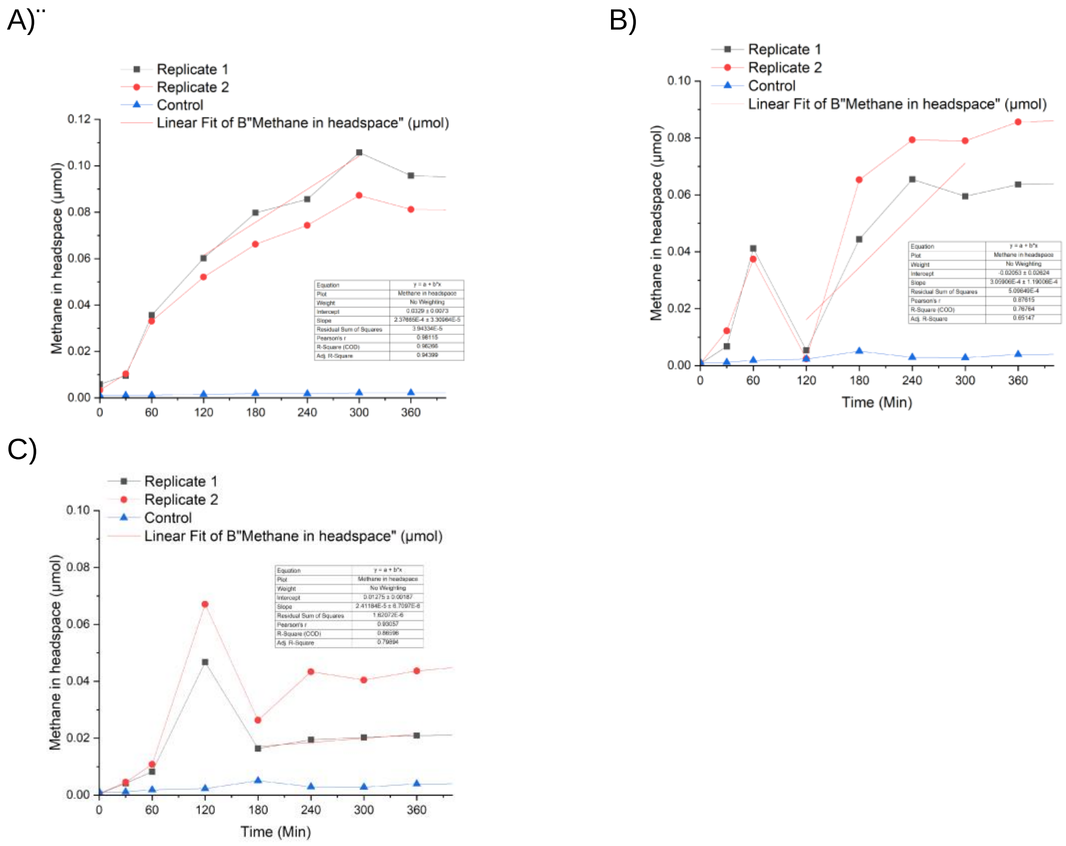


**Figure S13.** Linear regression of gas formation from alkyl-CoM mediated by Methanothermobacter marburgensis grown on $H_2/CO_2$ in bioreactor. Mcr assay was conducted at 65 °C using 1.1 mM vitamin $B_{12}$, 20 mM Ti(III) citrate, 0.5 mM of heterodisulfide of coenzyme B, and different concentration of alkyl-CoM as following: **(A)** 5 mM of methyl-CoM; **(B)** 4.5 mM of methyl-CoM & 0.5 mM of ethyl-CoM; **(C)** 2.5 mM of methyl-CoM & 2.5 mM of ethyl-CoM; **(D)** & **(E)** 0.5 mM of methyl-CoM & 4.5 mM of ethyl-CoM; **(F)** & **(G)** 5 mM of ethyl-CoM. All of the experiments were conducted with 100 µg of cell-free lysate from *Mt. marburgensis.*

D)

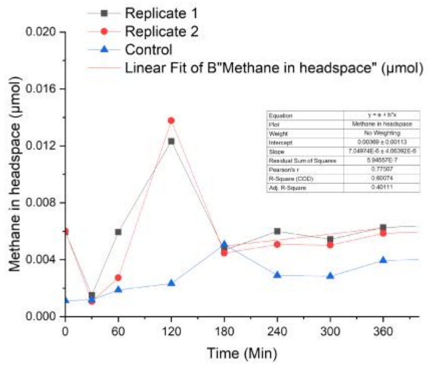


E)

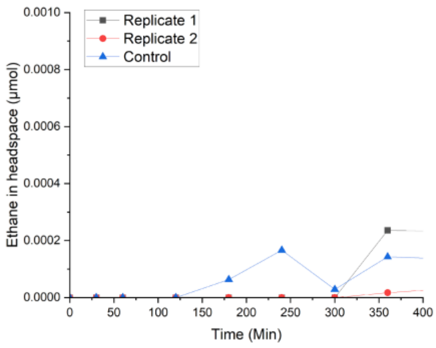


F)

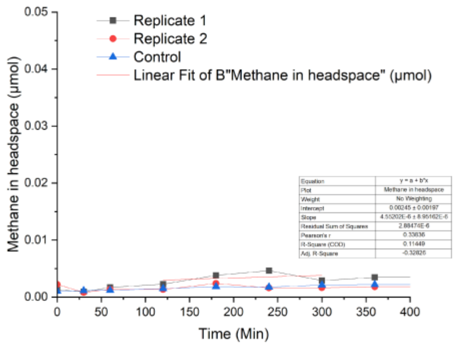


G)

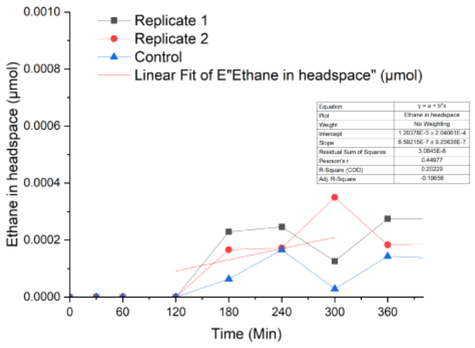


**Figure S14**. (continued)

A

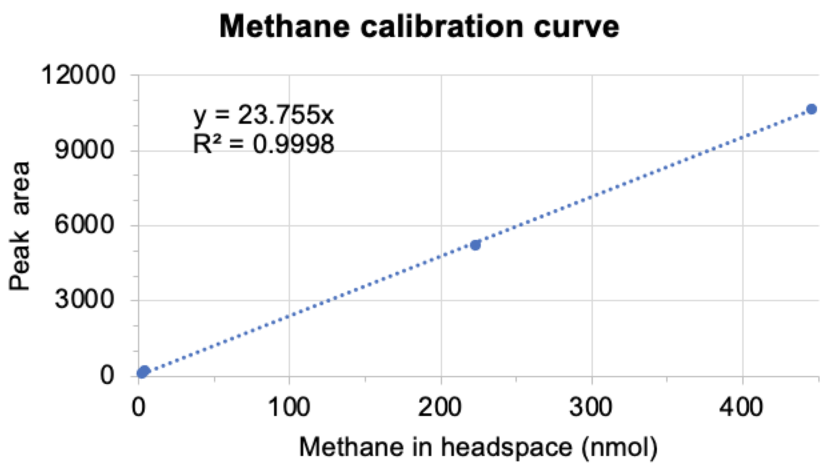


B

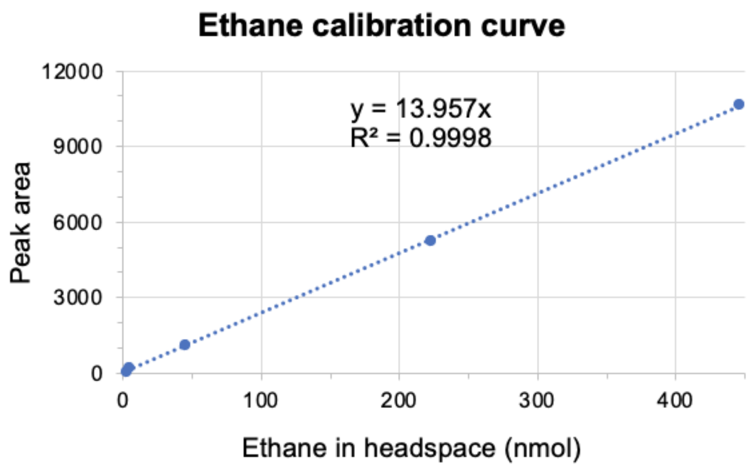


**Figure S15.** Methane and ethane calibration curves

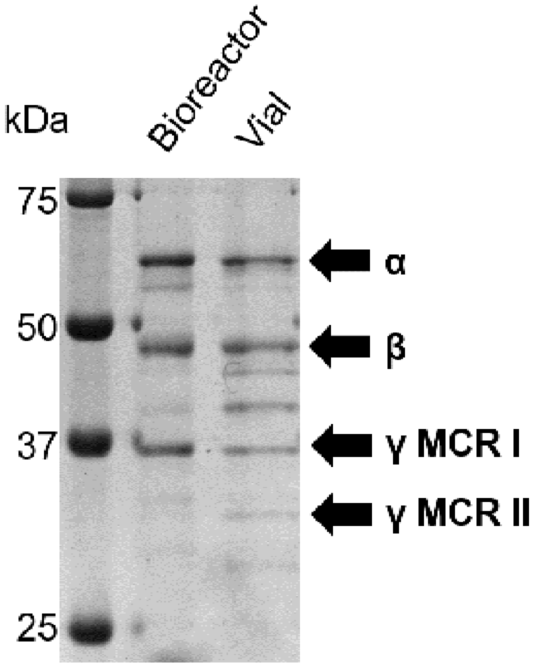


**Figure S16.** SDS-PAGE of ammonium sulfate-precipitated *M. marburgensis* Mcr extracts collected from two different growth conditions. The bioreactor and the vial sample samples were harvested at high- and low-optical density, respectively. The sizes of the subunits of both Mcr isoforms are: 65 kDa (alpha), 49 kDa (beta), 38 kDa (gamma of Mcr I), and 33 kDa (gamma of Mcr II).
